# *speck*, first identified in *Drosophila melanogaster* in 1910, is encoded by the Arylalkalamine N-acetyltransferase (AANAT1) gene

**DOI:** 10.1101/800391

**Authors:** Eric P. Spana, Amanda B. Abrams, Katharine T. Ellis, Jason C. Klein, Brandon T. Ruderman, Alvin H. Shi, Daniel Zhu, Andrea Stewart, Susan May

## Abstract

The pigmentation mutation *speck* is a commonly used recombination marker characterized by a darkly pigmented region at the wing hinge. Identified in 1910 by Thomas Hunt Morgan, *speck* was characterized by Sturtevant as the most “workable” mutant in the rightmost region of the second chromosome and eventually localized to 2-107.0 and 60C1-2. Though the first *speck* mutation was isolated over 115 years ago, *speck* is still not associated with any gene. Here, as part of an undergraduate-led research effort, we show that *speck* is encoded by the *Arylalkylamine N-acetyltransferase 1* (*AANAT1*) gene. Both alleles from the Morgan lab contain a retrotransposon in exon 1 of the RB transcript of the *AANAT1* gene. We have also identified a new insertion allele and generated multiple deletion alleles in *AANAT1* that all give a strong *speck* phenotype. In addition, expression of *AANAT1* RNAi constructs either ubiquitously or in the dorsal portion of the developing wing generates a similar *speck* phenotype. We find that *speck* alleles have additional phenotypes, including ectopic pigmentation in the posterior pupal case, leg joints, cuticular sutures and overall body color. We propose that the acetylated dopamine generated by *AANAT1* decreases the dopamine pool available for melanin production. When *AANAT1* function is decreased, the excess dopamine enters the melanin pathway to generate the *speck* phenotype.

## INTRODUCTION

In the early 20^th^ century, Thomas Hunt Morgan, Calvin Bridges, and colleagues began to identify mutations in *Drosophila* that produced visible phenotypes in adults. Some of these mutations have led to groundbreaking discoveries: *white* mutants verified the chromosomal theory of inheritance (Morgan, 1910), *Notch* mutants (Dexter, 1914; Morgan & Bridges, 1916) lead to the discovery of an oncogenic signal transduction pathway and a mechanism for lateral inhibition in cell fate determination in development (Siebel & Lendahl, 2017); and *Ultrabithorax* (or *bithorax* as it was known then (Bridges, C. B. & Morgan, 1923)) became a founding member of the homeobox family and an important factor in anterior/posterior patterning in metazoan development (Pick, 2016).

Remarkably, some of Morgan and colleagues’ mutant strains have only recently been assigned to a gene, while others still await identification. Many of these mutants fall into a few phenotypic categories like wing shape or bristle morphology or pigmentation (in eyes or body). In recent years, the *Curly* mutation (Ward, 1923) was identified as an allele of the *Duox* gene, though it was identified almost 100 years prior (Hurd, Liang, & Lehmann, 2015). The *straw* mutation (Morgan, Bridges, & Sturtevant, 1925) was mapped to the *laccase2* gene exactly 100 years after it was discovered (Sickmann, Gohl, Affolter, & Muller, 2017). In addition, a recent systematic screen has mapped many unannotated mutations, in some cases, to single genes (Kahsai & Cook, 2018).

In *Drosophila*, pigmentation of the adult cuticle is a combination of melanin and sclerotin which are end-products of branches of a shared biosynthetic pathway (Figure 1, reviewed in (Yamamoto & Seto, 2014) and (Massey & Wittkopp, 2016)). The pigmentation pathway begins with the sequential modification of L-tyrosine to L-Dopa by Tyrosine Hydroxylase (encoded by the *pale* gene) and to Dopamine by Dopa Decarboxylase (encoded by the *Ddc* gene). At dopamine, the pathway branches onto three routes: the melanin pathway, the NBAD sclerotin pathway and the NADA sclerotin pathway. Both L-Dopa and dopamine function as precursors for the extracellular production of melanin through quinone intermediates produced by the multicopper oxidase activity of the *straw* (*laccase2*) gene (Riedel, Vorkel, & Eaton, 2011). The *yellow* gene is required to produce melanin in the extracellular space but its exact activity in the process is not known (Hinaux et al., 2018) though paralogs *yellow-f* and *yellow-f2* are dopachrome isomerases (Han et al., 2002); one of the last steps in melanin synthesis. In the NBAD sclerotin pathway, the enzyme NBAD synthase (encoded by the *ebony* gene) combines the dopamine with beta-alanine and the product, N-Beta-alanyldopamine (NBAD) is secreted into the extracellular space where it is also converted to a quinone by *straw*. The beta-alanine is the product of the Aspartate decarboxylase enzyme (encoded by the *black* gene). Loss of *ebony* or *black* allows more dopamine to enter the melanization pathway causing a dark body color. Finally, dopamine may be converted to N-acetyldopamine (NADA) by the activity of Arylalkylamine N-acetyltransferase 1 (AANAT1) (previously called Dopamine Acetyltransferase, Dat). NADA secreted into the extracellular space may also converted to a quinone by *straw* and used in the formation of NADA sclerotin, which is colorless.

**Figure 1:**
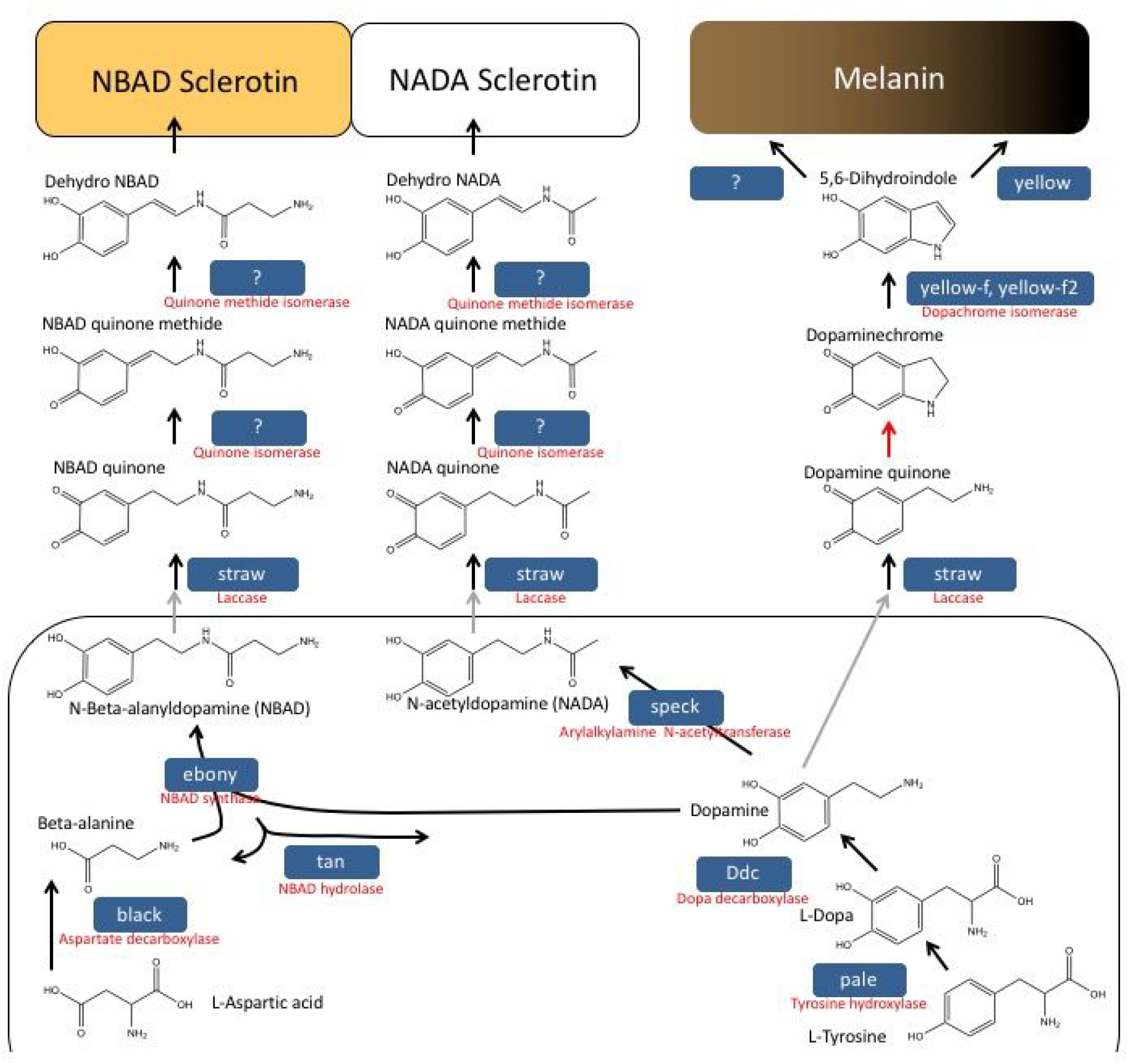
The pigmentation pathway in *Drosophila*. The biochemical pathway for the creation of NBAD Sclerotin, NADA Sclerotin and Melanin from L-Tyrosine and L-Aspartic acid is shown. The intercellular components are shown toward the bottom and exported from the cell (grey arrows) into the extracellular space. Black arrows indicate enzymatic reactions and red arrows are thought to be spontaneous reactions. *Drosophila melanogaster* gene names are shown in the blue boxes, and their enzymatic names are shown in red below them. Some enzymatic functions have been identified, but not associated with a gene.

The *Drosophila melanogaster* AANAT1 enzyme is a member of the GCN5-related N-acetyltransferase (GNAT) superfamily and transfers an acetyl group from acetyl-CoA to biogenic amines including dopamine, serotonin and others via a sequential binding mechanism (Cheng, Liao, & Lyu, 2012; Dempsey, D. R. et al., 2014). The *AANAT1* locus has two splice isoforms (RA and RB) and is expressed throughout development in a subset of gut and neuronal cells (Brodbeck et al., 1998; Hintermann, Jeno, & Meyer, 1995; Hintermann, Grieder, Amherd, Brodbeck, & Meyer, 1996). A partial loss of function allele (*AANAT1^lo^*), which shows decreased levels of isoform RA, shows no pigmentation phenotype but has behavioral defects in sleep recovery (Ganguly-Fitzgerald, Donlea, & Shaw, 2006; Shaw, Cirelli, Greenspan, & Tononi, 2000). Strong loss of function phenotypes in *Tribolium* (flour beetle) (Noh, Koo, Kramer, Muthukrishnan, & Arakane, 2016), *Bombyx* (silk moth) (Dai et al., 2010; Zhan et al., 2010), *Zootermopsis* (termite) (Masuoka & Maekawa, 2016), and *Oncopeltus* (milkweed bug) (Liu, Lemonds, Marden, & Popadić, 2016) show increased melanization and *AANAT1* has also been identified as the locus responsible for interspecies pigment differences in *Drosophila* (Ahmed-Braimah & Sweigart, 2015).

In a project initiated in an undergraduate laboratory course, we have mapped the *speck* mutation using genetic complementation and identified lesions in the gene encoding *Arylalkylamine N-acetyltransferase 1* (*AANAT1*). Both *speck^1^* and *speck^2^* contain the same 412 retrotransposon, though the *speck^2^* phenotype is stronger than *speck^1^*. We have generated new deletion alleles that show a darkening of the adult cuticle and additional phenotypes. All of these phenotypes can be phenocopied by RNAi and rescued by expression of a transgene. Loss of the *AANAT1* activity would allow more dopamine to enter the melanin biosynthesis and/or NBAD sclerotin biosynthesis pathway and produce darker animals in a mechanism similar to other insects.

## RESULTS

Though undergraduate scientists have been a staple of *Drosophila* research groups since Alfred Sturtevant worked for Morgan, only relatively recently has this research been extended to the classroom. A multi-year project in lab courses at the University of California, Los Angeles identified mutations responsible for eye development (Call et al., 2007) and a multi-year, multi-University consortium directed from Washington University in St. Louis had over 900 undergraduate students finish and annotate the dot chromosomes of a number of *Drosophila* species (Leung et al., 2015). Other <u>C</u>ourse-based <u>U</u>ndergraduate <u>R</u>esearch <u>E</u>xperiences (CUREs) have gained more prominence at Universities as faculty identify the advantages of such an approach (Shaffer et al., 2014) (see CUREnet: curenet.cns.utexas.edu for additional information) and using genetic complementation to characterize mutants derived from a screen has been quite successful (Bieser et al., 2019; Stamm et al., 2019). The success at the University level has led to this approach moving into high schools, where one collaboration between a high school and University yielded a LexA *Drosophila* enhancer trap collection (Kockel et al., 2016).

The creation of a molecularly defined deficiency kit (Cook et al., 2012; Roote & Russell, 2012) and two separate X chromosome duplications sets (Cook et al., 2010; Venken et al., 2010) allowed the simple, rapid mapping of *Drosophila melanogaster* mutations to a precise genomic interval by complementation. In combination with the ongoing transposable element mutagenesis from the Gene Disruption Project (Bellen et al., 2011), we hypothesized that undergraduate students could map mutations with adult visible phenotypes to a single gene within a semester. To decide which mutations to map first, we took a historical perspective and chose the four oldest mutations that FlyBase designated as unannotated (not associated to a transcription unit), that still maintained the adult, visible phenotype as described, and had stocks available: *speck* (*speck*, first identified in 1910), *curved* (*cu*, 1911), *spread*, (*sprd*, 1913), and *tilt* (*tt*, 1915). We found that *cu^1^* contained a ∼7 kb 412 Element retrotransposon (position 15,953,913, Genome annotation release R6.27) in an exon of the Stretchin-Mlck (Strn-Mlck) gene as described in (Rodriguez, 2004) and failed to complement an insertion in Strn-Mlck (pBac{PB}Strn-Mlck^c02860^). We found that the *spread* stock died prior to 1923 (Bridges & Morgan, 1923) and that all stocks listed as *sprd^1^* are likely mis-annotated. We mapped *tt^1^* to a defined molecular region and *speck* to a single gene, which we further characterized outside of class.

### History of the *speck* mutation

Thomas Hunt Morgan isolated the first *speck* mutation in March 1910. However, that first *speck* mutant “was set aside in order that more time might be given to the study of the sex-linked eye-color *white* which had appeared in April 1910” (Bridges, C. B. & Morgan, 1919). A second, phenotypically stronger allele of *speck* was identified by Morgan in May 1911 (occasionally referred to as *olive speck* in their writing due to its slightly darker body color), and unfortunately, the 1910 stock was discarded. This 1911 *olive speck* mutation is what is currently known as *speck^1^*. A third *speck* allele was identified in June 1925 by Calvin Bridges from a stock (*al b pr cn vg a speck^1^*) that had an unusually dark body color than would be expected of a *black speck* double mutant. Subsequent crosses and tests led Bridges to conclude that the chromosome carried a new, stronger *speck* allele, which he named *speck^2^* (Bridges, Calvin B., Unpublished Material). The *speck* phenotype was described as a dark, melanized region at the hinge (axil) of the wing and a slight darkening of the body color. Bridges also noted “darker, brooch-shaped” region at the former anal region of the pupal case. As the adult phenotype was easy to score and *speck* mapped to the distal region of the right arm of chromosome 2 (2-107.0), *speck* has been a useful tool for genetic recombination mapping since the early 20^th^ century. Because *speck^2^* adults have a slightly darker body color, *speck* was predicted to function in the melanization or sclerotinization pathway (Wright, 1987).

### Mapping the *speck* mutation and generating new alleles

The easiest *speck* phenotype to score is the melanized region at the axil of the wing (Bridges & Morgan, 1919) (Figure 3A and B) and we chose this as the phenotype to score in complementation tests. At the start of the course there were two available alleles of *speck*: *speck^1^* and *speck^2^*; and *speck^2^* is also found on the SM5 and SM6 balancer chromosomes. Previous mapping had placed *speck* in the cytological region of 60C1,2; which roughly corresponds to more than 60 kb of DNA containing approximately 12 protein coding transcripts (Peng & Mount, 1990). To account for the variance in breakpoints between cytologically defined deficiencies and molecularly defined deficiencies, we tested the complementation of both *speck^1^* and *speck^2^* with six deficiency strains that covered cytological region 60B8 to 60D14 (Figure 2A). We found that two of the six deficiency strains failed to complement *speck* (*Df(2R)BSC356* and *Df(2R)BSC155*), molecularly defining the *speck* interval as between the left breakpoint of *Df(2R)BSC155* (by inclusion) and the left breakpoint of *Df(2R)BSC780* (by exclusion), a region containing approximately 26 protein coding transcripts (Figure 2B).

**Figure 2:**
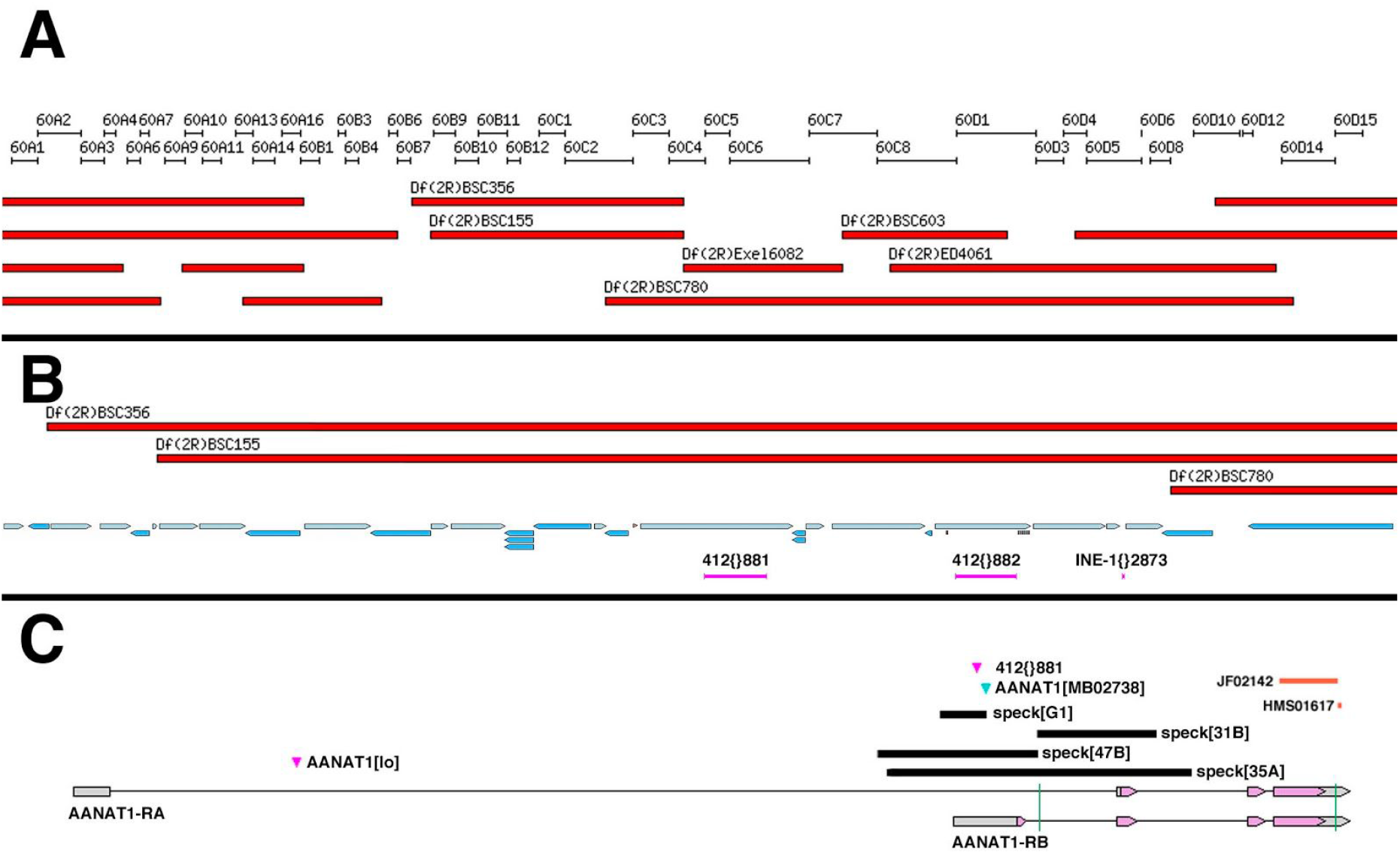
The *speck* mutations are lesions in the *AANAT1* locus. A.) A 900 kb region of the second chromosome encompassing cytological bands 60A1 through 60F15 shows the position of the deficiency strains used for complementation tests with *speck*. Only *Df(2R)BSC356* and *Df(2R)BSC155* failed to complement *speck*. B.) A 172 kb region that includes 60B7 through 60C2 shows the 21 transcription units (blue arrows) that lie within the left breakpoint of *Df(2R)BSC155* and the left breakpoint of *Df(2R)BSC780*. Within this region lay two 412 retrotransposons and an INE element in the *Drosophila* genome reference strain (*y^1^; cn^1^ bw^1^ speck^1^*). C.) A map of the *AANAT1* locus as it appears in a wild type genome. The 412{}881 element is only found in the *speck^1^* and *speck^2^* alleles and not in wild type. The *AANAT1^MB02738^* insertion and the small deletion (black bar) generated from it, *speck^G1^*, are also *speck* alleles. The deleted regions of the CRISPR derived alleles, *speck^31B^*, *speck^47B^* and *speck^35A^* are shown with black bars. The locations of the two RNAi regions (JF02142 and HMS01617) are located at the 3’ end of the transcript and can also produce a *speck* phenotype. The vertical green lines through the transcripts show the approximate locations of the gRNAs used to make the CRISPR alleles.

**Figure 3:**
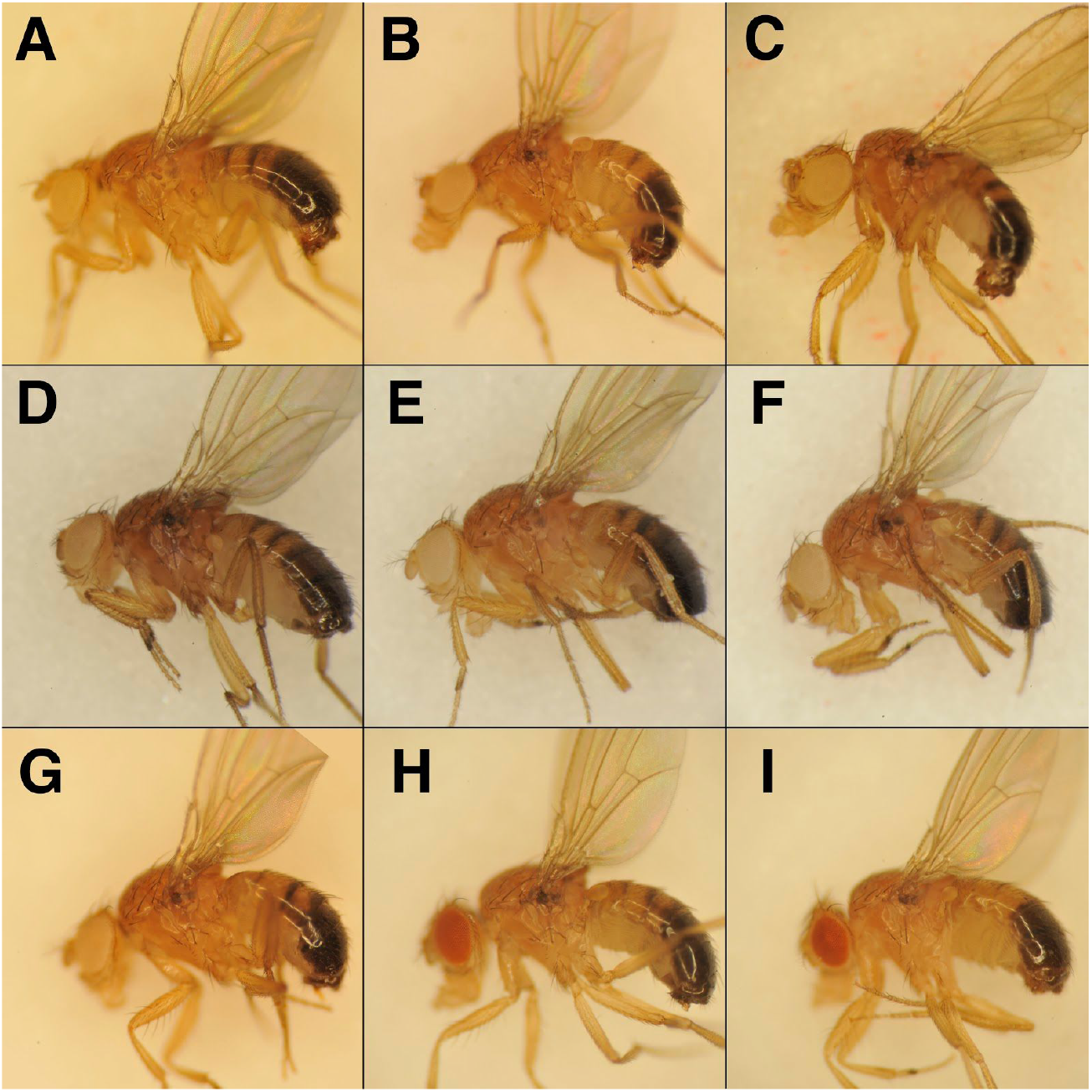
Mutations in the *AANAT1* gene cause the *speck* phenotype of ectopic pigmentation in the wing hinge and darker body color. Adult males are shown with anterior to the left and dorsal up. A.) A *w^1118^* male shows the wild type pigmentation at the ventral wing hinge region. B.) The *speck^1^* allele (*cn^1^ bw^1^ speck^1^*) shows a “speck” of pigmentation at the ventral wing hinge. C.) The speck^2^/Df genotype (*Df(2R)BSC356*/SM6a) shows the dark speck at the wing hinge and a slightly darker body color. D.) An insertion allele of *AANAT1* (*Mi{ET1}AANAT1^MB02738^*) shows a darker body color and a pronounced wing hinge spot. E.) Mobilization of the insertion in D can revert the *speck* phenotype (*speck^T1^*) F.) An imprecise excision of the insertion in D that removes an *AANAT1* transcriptional start site produces a *speck* phenotype (*speck^G1^*). G.) A CRISPR allele (*speck^47B^*) that deletes 1218 bp shows an intermediate *speck* phenotype. H. & I.) Knock-down of the *AANAT1* gene by two different GAL4 drivers and two different hairpin locations produce a *speck* phenotype (H: Tubulin-GAL4, UAS-dicer2; UAS-AANAT1^HMS01617^ and I: apterous-GAL4; UAS-AANAT1^JF02142^).

Genetic interactions with *speck* helped to further narrow the list of candidate genes. The *suppressor of sable*, *su(sable)* mutation, suppresses aspects of the *speck^1^* phenotype (Warner, Grell, & Jacobson, 1975) and *su(sable)* also suppresses phenotypes caused by 412 or roo retrotransposon insertions in a number of genes. *In situ* hybridization showed a 412 element was present in the genomic region near *speck* (Searles & Voelker, 1986), indicating that the *speck^1^* allele may be derived from a 412 insertion.

In a lucky coincidence, the *speck^1^* allele was sequenced by whole genome sequencing—it was present in the isogenic strain (*y^1^; cn^1^ bw^1^ speck^1^*) used for the Drosophila Genome Project (Adams et al., 2000) and is therefore displayed as the reference strain in the genome browsers in FlyBase. Two 412 elements are present in the region defined by the deficiency breakpoints: 412{}881 and 412{}882 (Figure 2B). The 412{}881 element is not present in any *speck^+^* strains (by PCR or analysis of whole genome sequencing). It is present in multiple stocks of both *speck^1^* and *speck^2^*. The 412{}881 element is inserted in the 5’ UTR in exon 1 of the gene encoding *Arylalkylamine N-acetyltransferase 1* (*AANAT1*) splice isoform RB. An allele of *AANAT1*, *AANAT1^lo^* (Figure 2C) has been shown to cause a decrease in the AANAT1-RA product, however we find that *AANAT1^lo^* fully complements all *speck* alleles. Figure 2C shows the AANAT locus as it appears in wild type genomes which matches the gene structure of the locus described previously (Brodbeck et al., 1998).

We attempted to identify the molecular difference between *speck^1^* and *speck^2^*. Whole genome sequencing of two different strains of *speck^2^* were sequenced as heterozygotes with the prediction that changes in the *AANAT1* locus that were shared between them would warrant further examination as being the *speck^2^* lesion. Assembly of the *AANAT1* loci from both and comparison to the s*peck^1^* reference strain did not yield any candidate changes (data not shown). In addition, examination of SNPs between balancers carrying *speck^2^* and the reference strain carrying *speck^1^* showed no SNPs within the AANAT1 locus (Miller et al., 2018). Because there were no changes in the genomic interval that short-read, whole genome sequencing could identify, we looked to see if 412{881} had a second retrotransposon insertion, similar to AANAT1^lo (Brodbeck et al., 1998)^. However, a PCR product spanning the 412{881} insertion was identical in size and structure in both s*peck^1^* and *speck^2^* (data not shown). Though *speck^2^* has a stronger phenotype (see below), we have not been able to identify any molecular change.

We obtained more definitive evidence that *speck* mutants disrupt *AANAT1* from the study of a strain containing a Minos insertion, AANAT1^MB02738^. This insertion is in the same exon as the 412 insertions in *speck^1^* and *speck^2^* and displays a strong *speck* phenotype (Figures 2C and 3D) and fails to complement *speck^1^* and *speck^2^* alleles. To test whether the Minos insertion causes the *speck* phenotype we mobilized the Minos insert with a hs-Minos transposase (SM6a, P[hs-MiTpase) (Metaxakis, Oehler, Klinakis, & Savakis, 2005) and recovered 10 unique revertant lines that lacked the transgenic marker associated with the Minos ET1 vector (GFP). Of these 10 GFP revertants, 4 maintained the *speck* phenotype as homozygotes, but 6 fully reverted the *speck* phenotype to wild type. One allele, *speck^T1^*, reverted the *speck* phenotype and had only a 6 nucleotide insertion remaining from the Minos element (Figure 3E). One of the Minos revertant alleles, *speck^G1^* was an imprecise excision that deleted over 500 bases, including the AANAT1-RB transcription start site (Figures 2C and 3F). In addition, we identified an unusual allele from this mobilization, as well. The *speck^D1^* allele (not shown) reverts the *speck* phenotype to wild type and has removed the entire Minos element, but has inserted approximately 480 bp of 412 element into the site where the Minos element had been. Presumably, the double stranded break created by the mobilization of the Minos element was repaired from the homologous chromosome, the SM6 balancer carrying the *speck^2^* allele, providing a nice example of gene conversion.

To make null alleles that would remove both *AANAT1* splice isoforms, we attempted to make deletion alleles using CRISPR (Gratz et al., 2013). We identified two Cas9 guide RNA binding sites: one in intron 1 of the AANAT1-RB isoform and another in the 3’ UTR shared by both isoforms. Two Cas9 cuts and non-homologous end-joining would create a deletion lacking all the functional domains of AANAT1. After injections of the two guide-RNA plasmids into Cas9 expressing embryos, we crossed individual surviving adults to the *w*; *AANAT1^MB02738^* stock and scored their progeny for a *speck* phenotype. Out of 66 crosses, we identified 3 CRISPR derived *speck* alleles and sequenced PCR products that spanned the lesions in each. The strongest allele *speck^35A^*, removes exons 1 and 2 of *AANAT1-RB* and exon 2 of *AANAT1-RA* (Figure 2C). There should be no protein generated from the *AANAT1-RB* isoform (since its transcription should never be initiated) and since exon 2 of *AANAT1-RA* has the translation initiation codon, creation of a protein product from *AANAT1-RA* in *speck^35A^* would necessitate an exon skipping event from exon 1 to 3, and for the first AUG in exon 3 to be in the correct frame (it is not). Like *AANAT1^G1^*, the CRISPR derived allele *speck^47B^* also removes only exon 1 of *AANAT1-RB* and shows a *speck* phenotype (Figure 2C and 3G). The *speck^31B^* CRISPR allele removes exon 2 from both *AANAT1* isoforms (Figure 2C) and shows a *speck* phenotype (data not shown).

Additionally, we found that we could replicate the *speck* phenotype using RNAi against the *AANAT1* gene. Two different *AANAT1* hairpin constructs under UAS control (Figure 2C) produced a *speck* phenotype using the drivers *Tubulin-GAL4*, which is expressed ubiquitously and *apterous-GAL4*, which is expressed in the dorsal region of developing wing tissue (Figure 3H and 3I) but not engrailed-GAL4, which is expressed in the posterior region of developing wing tissue (data not shown). One interpretation would be that the hinge tissue giving rise to the pigmented region is derived from dorsal, but not posterior tissue in the wing imaginal disc. In summary, through multiple genetic approaches we identified that *speck* mutants are derived from disruptions in the *AANAT1* gene.

### Phenotypic Characterization

To determine when the *speck* phenotype first presents, we performed time-lapse photography on newly eclosed, immobilized flies. During normal development (Supplemental Movie 1), wing expansion and sclerotization and melanin deposition (tanning) occurs during the first three hours after eclosion from the pupal case. In *speck^2^* (*speck^2^, bs^2^*) we find that the pigmented regions at the wing hinge are not evident at eclosion, but become more pronounced during the cuticle tanning and sclerotization period (Supplemental Movie 2). The wing blister phenotype is due to the *blistered* allele present in the stock.

The *speck^2^* allele was originally described as being darker than *speck^1^*. While we also see *speck^2^* alleles as slightly darker than *speck^1^*, we found that *AANAT1^MB02738^* and some deletion alleles are much darker and present additional phenotypes. We find that the *AANAT1^lo^* allele (Figure 4B) has no obvious ectopic pigmentation and looks quite similar to a wild type strain, *w^1118^* (Figure 4A). The *AANAT1^MB02738^* allele (Figure 4C) shows not only strong ectopic pigmentation in the wing hinge, but additional areas. The intersection of the scutellum and mesothoracic laterotergite shows a highly melanized spot (arrowhead in enlarged image) and the joints also show ectopic pigmentation (arrow). In addition, the sutures between cuticle plates on the lateral body wall are more pronounced, however this may be caused by the overall darkened cuticle. The CRISPR derived alleles *speck^31B^* and *speck^35A^* show a very similar phenotype to *AANAT1^MB02738^* (Figures 4D and 4E) with regard to body color, wing hinge, scutellum, and leg joint pigmentation. The *speck^47B^* allele, however, shows a phenotype more similar to *speck^1^*, showing only the wing hinge phenotype.

**Figure 4:**
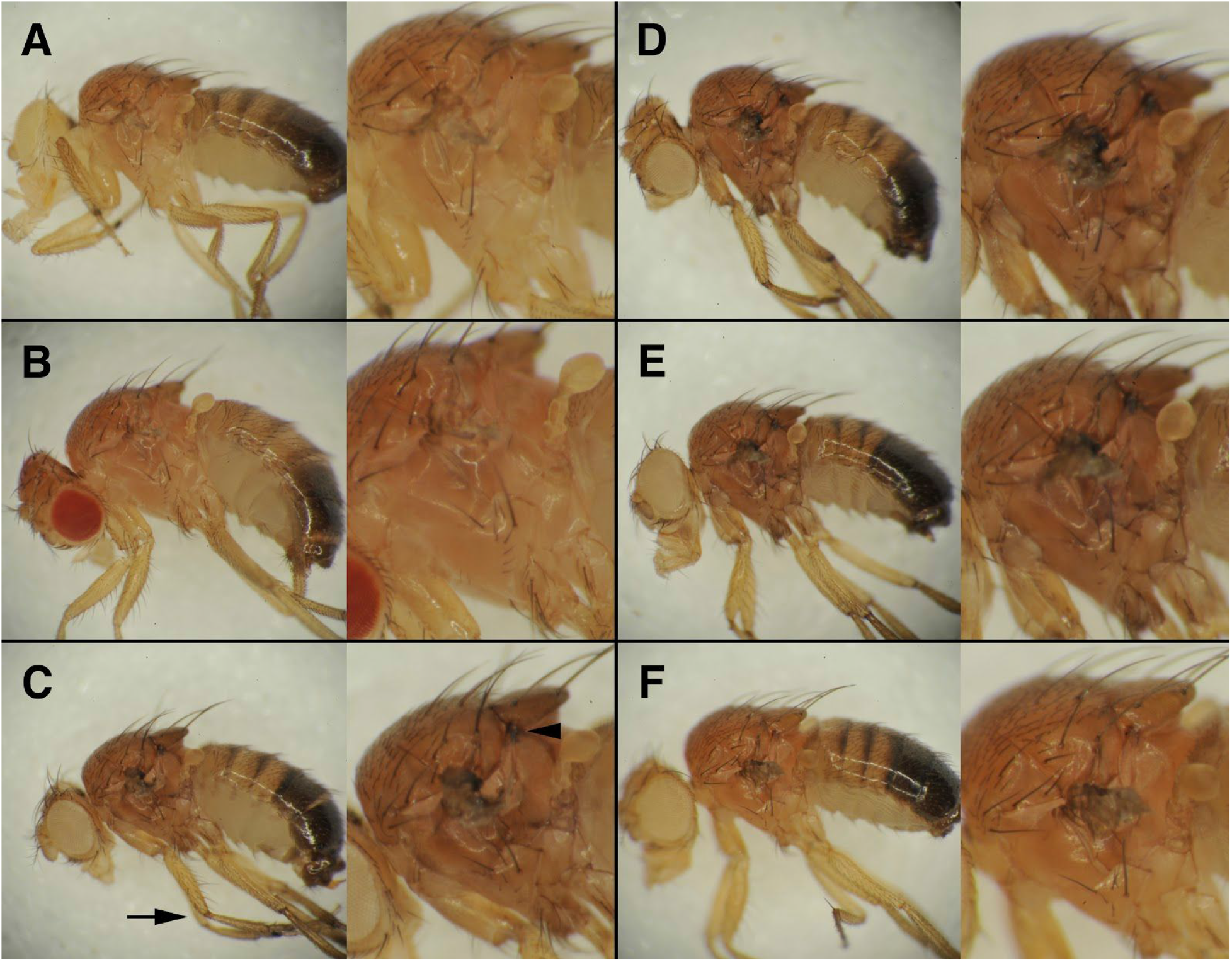
Strong *speck* alleles show pigment accumulation in leg joints and thoracic cuticle sutures. Seven-day-old males whose wings were removed are shown with anterior to the left and dorsal up. Each panel shows the whole fly on the left and a 2X zoom on the right. A.) A *w^1118^* male shows no ectopic pigment accumulation in the wing hinge or any other cuticular regions. B.) The *AANAT1^lo^* allele (*bw^1^ AANAT1^lo^*) is essentially wild type with regard to cuticle pigment. C.) Flies homozygous for the *AANAT1^MB02738^* insertion (*w^1118^; AANAT1^MB02738^*) show not only ectopic pigmentation at the wing hinge region, but at the region between the scutellum and the mesothoracic preepisternum (arrowhead), and also the leg joints (arrow). In addition, they show an overall darker pigment to the thorax, which makes the regions between portions of the body wall more evident. D and E.) Males of *w^1118^; speck^31B^* (D) and *w^1118^; speck^35A^* (E) show a similar phenotype to those in C. F.) *w^1118^; speck^47B^* males show only the wing hinge phenotype.

The wing hinge and body color are not the only described phenotypes of *speck* mutants. Both Bridges (Bridges, Unpublished Material), unpublished notes) and Waddington (Waddington, 1942) described a dark anal pad of the pupal case during pupariation. Because this phenotype had been described, but never shown in a publication, we documented the anal pad phenotype of *speck* mutants. We find that compared to wild type (Figure 5A) the anal pad in both *speck^1^* (Figure 5B) and *speck^2^* (Figure 5C) are substantially darker. This ectopic pigment is also present in *AANAT1^MB02738^* (Figure 5D) and can be reverted to wild type (Figure 5E) after mobilization of the Minos insertion. Furthermore, ubiquitous expression of a small hairpin against the *AANAT1* gene can phenocopy *speck* in the anal pad of pupae (Figure 5F).

**Figure 5:**
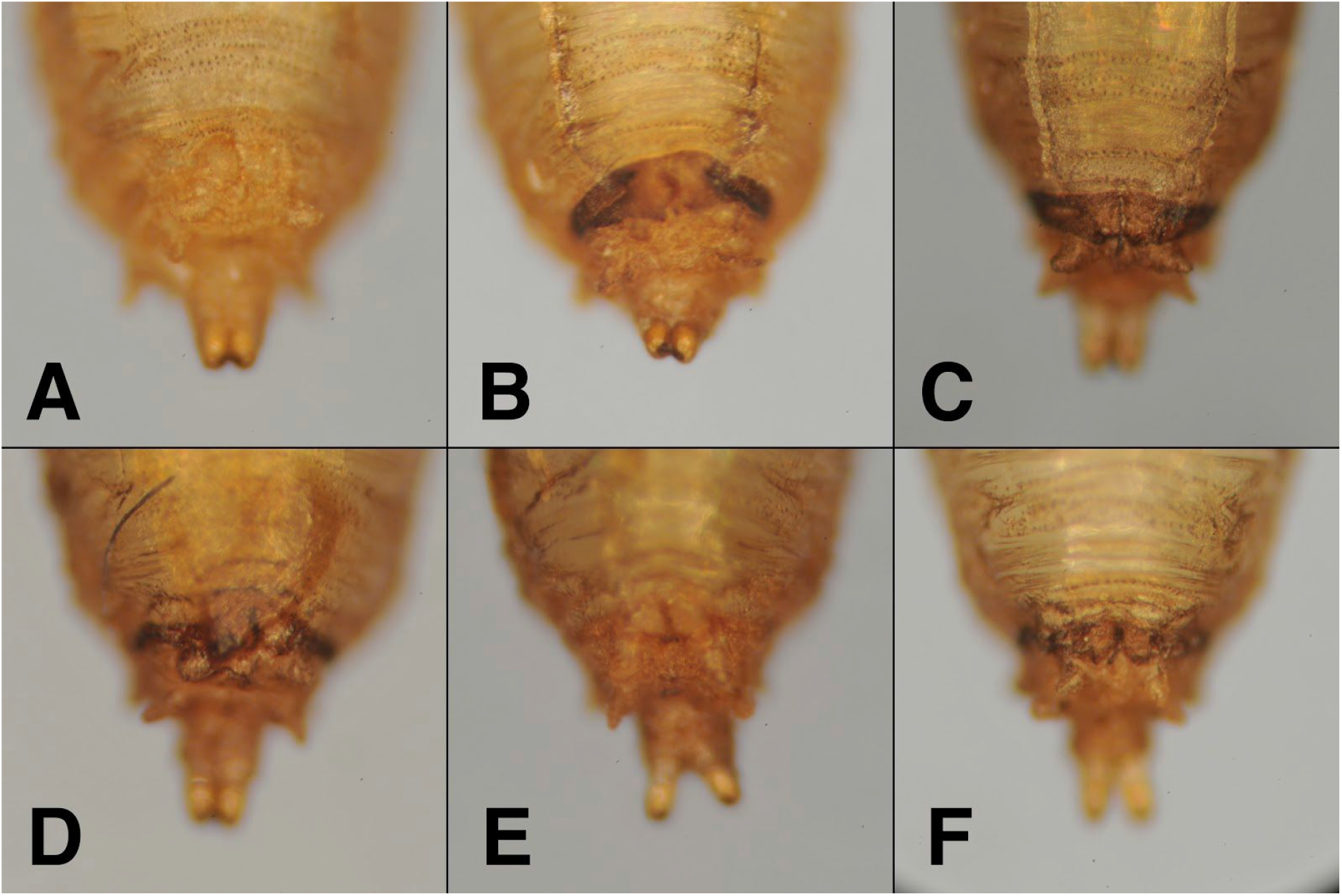
The pupal case of *speck* mutants display excess pigmentation at the anal pad. All images are of the posterior ventral sides of pupal cases following eclosion. A.) An Oregon R pupal case shows normal posterior pigmentation. B.) A *speck^1^* mutant (*cn^1^ bw^1^ speck^1^*) shows the anal pad region is darkly pigmented. C.) A *speck^2^* mutant (*speck^2^ bs^2^*) shows a similar dark band as *speck^1^* and the pupal case overall is slightly darker. D.) An insertion in *AANAT1* (*AANAT1^MB02738^*) shows a similar phenotype and can be reverted to wild type upon mobilization (E). F.) Knockdown of *AANAT1* via RNAi can reproduce the phenotype (Tubulin-GAL4, UAS-Dicer2; UAS-AANAT1^HMS01617^).

### Rescue and Ectopic Expression

We wanted to determine if expression of a related *AANAT* gene could rescue the pigmentation phenotype of *speck* mutants. Expression of the *Bombyx mori* ortholog of the *Drosophila AANAT1* gene has been shown to decrease pigment production in *Drosophila* when misexpressed via heat shock (Mizuko Osanai-futahashi et al., 2012). We used this transgenic construct (*UAS-Bm aaNAT*) to rescue the *speck* phenotype. Flies heterozygous for the *AANAT1^MB02738^* allele (*w^1118^; AANAT1^MB02738^/+*) display a wild type phenotype (Figure 6A) while *AANAT1^MB02738^/speck^2^* (*w^1118^; AANAT1^MB02738^/SM6a*) show a moderate *speck* phenotype (Figure 6B). Misexpression of *UAS-Bm aaNAT* using Tubulin GAL4 shows a stark decrease in pigment production (Figure 6C) (*w^1118^; AANAT1^MB02738^/+; Tubulin GAL4,* UAS-Bm aaNAT*/+*) and this loss of pigment is still evident in a *speck* background (*w^1118^; AANAT1^MB02738^/SM6a; Tubulin GAL4,* UAS-BM aaNAT*/+*) as seen in Figure 6D. However, the phenotype of wing hinge pigmentation is completely rescued. This loss of the *speck* phenotype in Figure 6D is not simply because the flies are lighter, as the *speck* phenotype is still obvious in a *yellow* background (Figure 6E). This rescue experiment, coupled with our sequencing and genetic mutant analysis verifies that *speck* mutations disrupt the *AANAT1* gene.

**Figure 6:**
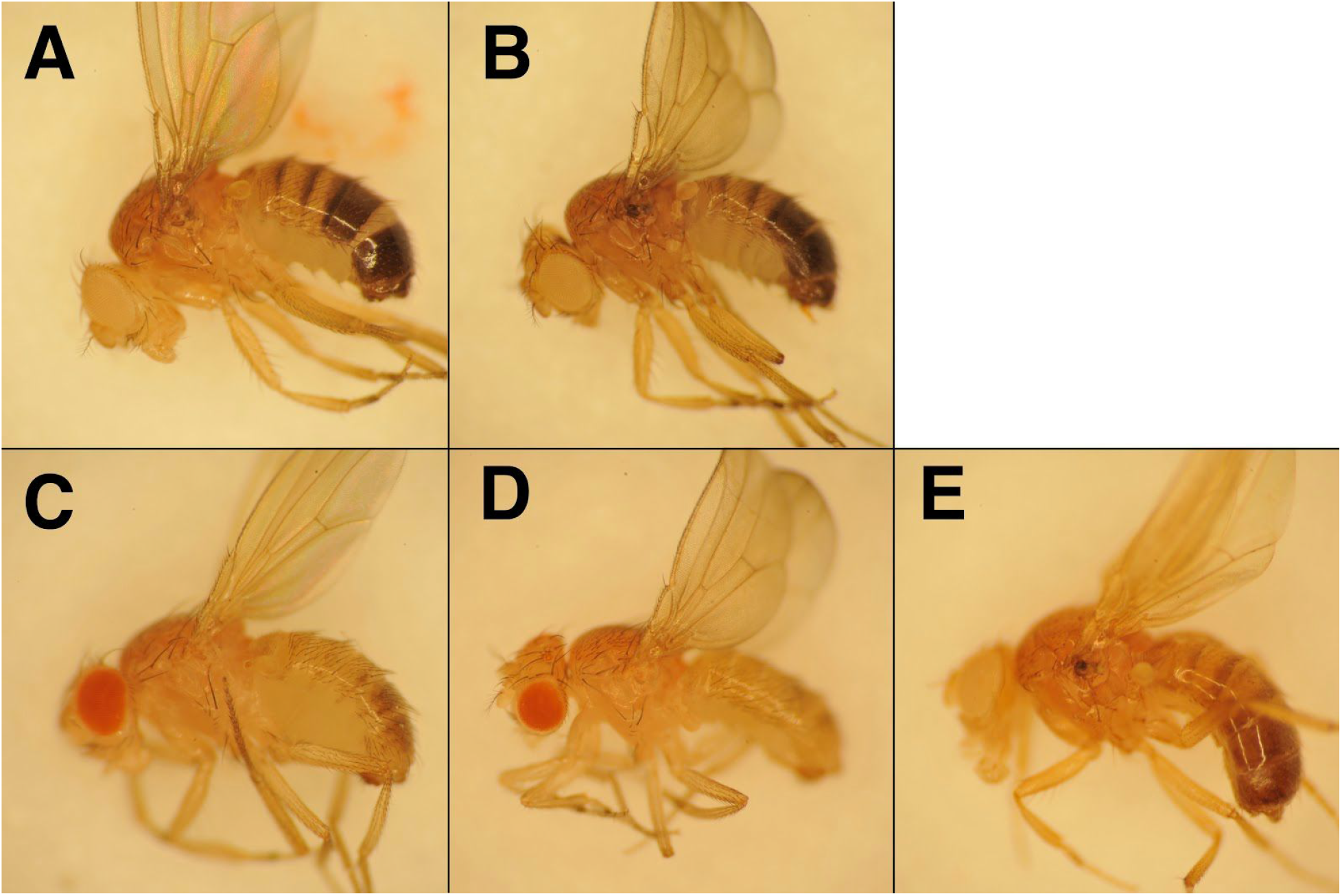
The *Bombyx mori* aaNAT gene can rescue *speck* mutations, but causes decreased pigmentation. Adult males are shown with anterior to the left and dorsal up. A.) A male heterozygous for the *AANAT1^MB02738^* allele shows wild type pigmentation (*w^1118^; AANAT1^MB02738^/+*). B.) The *speck* phenotype is observed in the *AANAT1^MB02738^/speck^2^* genotype (*w^1118^; AANAT1^MB02738^/SM6a*) C.) Ubiquitous misexpression of the *Bombyx mori* aaNAT gene causes a loss of pigmentation (*w^1118^; AANAT1^MB02738^/+*; TubulinGAL4, UAS-Bm-aaNAT/+) D.) Ubiquitous misexpression of the *Bombyx mori* aaNAT rescues the *speck* phenotype (*w^1118^; AANAT1^MB02738^*/SM6a; TubulinGAL4, UAS-Bm-aaNAT/+). E.)The *speck* phenotype in the wing hinge is still visible in genotypes that reduce pigmentation, such as *yellow*. (*y*w*; bw^1^, speck^1^, zip^2^*/SM6a)

## DISCUSSION

We have mapped the *speck* mutation, first identified by Thomas Hunt Morgan in 1910, to the *AANAT1* gene and found that both Morgan’s (*speck^1^*) and Bridges’ (*speck^2^*) alleles are retrotransposon insertions in the 5’ UTR of one of *AANAT1*’s two transcripts. However, no obvious differences in sequence between their two alleles were identified. A transposon mutagenesis screen almost 100 years after *speck* was discovered produced a Minos insertion in the same region that fails to complement *speck* and gives a similar phenotype. Excision of that Minos element can revert the *speck* phenotype, indicating that the insertion causes the phenotype. Deletion alleles that remove parts of both transcription units also give a *speck* phenotype indicating that *AANAT1* null flies are adult viable, fertile and have a *speck* pigmentation phenotype. Ubiquitous expression of the *AANAT1* ortholog of *Bombyx mori* can rescue the *speck* phenotype and also decrease the cuticular pigmentation. Because we find mutagenic insertions in *AANAT1* only in *speck* mutations, that removal of the insertions can revert the phenotype to wild type, that deletions of *AANAT1* produce a *speck* phenotype and that the phenotype can be rescued by a transgene, we can conclude that *speck* is a loss of *AANAT1*; that *AANAT1* is the *speck* gene.

### How does *speck* fit into the *Drosophila* cuticular pigmentation pathway?

While *AANAT1* has been known in the cuticular pigmentation pathway for some time, the *speck* phenotype is not what would have been predicted from a mutation that causes increases in melanin and NBAD sclerotin. As seen in Figure 1, *ebony* mutants would have a loss of tan pigment due to the loss of NBAD sclerotin, but increases of black/brown melanin and colorless NADA sclerotin due to dopamine being shunted into those pathways. One would predict that *speck* mutants would lose no visible pigments from loss of NADA sclerotin, but would have increases of black/brown melanin and tan NBAD sclerotin pigment. So how could *speck* mutants be lighter in body color than *ebony* mutants?

Since *ebony* (and *black*) mutants are much darker than *speck* mutants, there may be differences in the amount of NBAD versus NADA sclerotin in different parts of the cuticle. Presuming both *ebony* and *speck* mutants have the same amount of L-dopa and dopamine going into the choice points as reviewed in Figure 1, the strengths of the pathways could be different. If the dorsal thorax utilized the NBAD sclerotin pathway much more than the NADA sclerotin pathway, loss of *ebony* would send a substantial amount of precursors into the NADA pathway (which is colorless) and black/brown melanin pathways giving the *ebony* mutants their namesake color. So *ebony* mutants are very dark because they have moved enough dopamine out of the NBAD sclerotin pathway and into the melanin pathway to make very dark flies. However, if the amount of NADA sclerotin in the dorsal thorax was low, loss of *speck* would provide less dopamine for the NBAD and melanin pathways. So *speck* mutants are a little darker but not as dark as *ebony*. In other regions of the adult cuticle, like the wing hinge and leg joints, this might be reversed, implying those regions have higher concentrations of NADA sclerotin than NBAD sclerotin and have more precursors to send to the pathways yielding black/brown and tan pigments from increased melanin and NBAD sclerotin.

What might these differences reveal? The most obvious difference is that the dorsal cuticle is hard and rigid and the wing hinge and leg joints are soft and pliable. Might the ratio of NBAD versus NADA sclerotin make a difference in cuticle hardness? Drosophila *black* mutants show a significant increase in puncture resistance when injected with beta-alanine to rescue the *black* phenotype (Jacobs, 1985). A *Blattella* (cockroach) strain that had decreased amounts of beta-alanine also showed decreased puncture resistance (Czapla, Hopkins, & Kramer, 1990) and a similar result was found with *Tribolium* (Roseland, Kramer, & Hopkins, 1987). Though this is an interesting correlation, many factors play a part in cuticular hardness such as thickness, protein and chitin composition, and others (reviewed in (Andersen, 2012).

Another possibility, though is redundancy. Similar to *yellow* that has 12 *yellow*-related genes in the *Drosophila* genome (Drapeau, 2001), there are seven AANAT-like genes in the *Drosophila* genome in addition to *AANAT1* (Amherd, Hintermann, Walz, Affolter, & Meyer, 2000). It is possible that one or more of these genes could also function in sclerotization, and that *AANAT1* might not act alone. One of these AANAT-like genes, *AANATL2*, has been shown to be present in adult flies and catalyze the formation of long-chain acylserotonins and acyldopamines (Dempsey, Daniel R. et al., 2014), however this seems unlikely to function in the NADA sclerotin pathway.

### How does the speck Drosophila phenotype differ from other insects?

We find that strong *speck* alleles in *Drosophila* have ectopic melanin formation in regional positions and a slightly darker cuticle in adults. These locations in adults coincide with positions in the exoskeleton where flexibility is required, such as leg joints, wing hinges and suture locations between cuticle plates. Only the strongest of these phenotypes can be replicated by RNAi. In silkworms (*Bombyx mori*), mutations show an overall body darkening, and regional melanization in the larval head, thoracic legs and anal plate (Dai et al., 2010; Zhan et al., 2010). In adult milkweed bugs (*Oncopeltus*), no change in adult cuticle pigmentation was detectable by RNAi, though ectopic melanization was found in regions of the proximal hindwing (Liu et al., 2016). Expression analysis showed that region was the only position where AANAT was expressed in either the forewing or hindwing. Reduction of AANAT by RNAi in American cockroaches (*Periplaneta*) showed no forewing or hindwing phenotypes, but did show increased melanization in an unpigmented region of the T1 plate (Liu et al., 2016). In termites (*Zootermopsis*), reduction of AANAT by RNAi caused intense melanization of the anterior head and mandibles (Masuoka & Maekawa, 2016). Red flour beetles (*Tribolium*) injected with RNAi against AANAT showed a combination of phenotypes: increased melanization in adult body wall cuticle, elytra and hindwings (Noh et al., 2016).

Common phenotypes seen across these insect species fall into two categories: broad overall darkening of the adult cuticle and regions of intense melanin localization. While some phenotypic differences can be explained the variability of RNAi phenotypes and species specific development, some commonalities are striking. The loss of function mutations in *Drosophila* and *Bombyx* both caused overall body darkening and more intense pigmentation in legs and anal plates. Ectopic pigmentation was found in the proximal, posterior region of the hindwing in both *Oncopeltus* and *Tribolium*. It will be interesting to see how the expression pattern of *AANAT1* during cuticular tanning in *Drosophila* relates to its pigmentation phenotype.

### Possible defects in neuronal signaling in speck mutants?

In *Drosophila*, nervous system expression of *AANAT1* is typically associated with the inactivation of monoamine signaling by acetylation. In vertebrates, dopamine signaling is inactivated by a class of enzymes called monoamine oxidases (MAOs) that have, so far, not been identified in insects (reviewed in (Yamamoto & Seto, 2014)). As *AANAT1* is a cytoplasmic enzyme, inactivation of dopamine signaling, for example, would need to happen after endocytic uptake of synaptic dopamine by either the presynaptic or post-synaptic neuron since *AANAT1* expression has been described as neuronal, not glial. With the publication of the adult *Drosophila* brain “connectome” (Zheng et al., 2018), the identification of which synapses inactivate biogenic amines via *AANAT1* acetylation may soon be deduced. Weak alleles of *AANAT1* show a defect in sleep (Ganguly-Fitzgerald et al., 2006; Shaw et al., 2000), but it is not known how its expression pattern relates to the emerging network of sleep in *Drosophila* (Bringmann, 2018).

*AANAT1* might affect sleep by more than inactivating biogenic amines. In vertebrates, *AANAT1* orthologs have been termed a “Timezyme” ((Klein, 2007)) because its rhythmic night-time expression and/or activity are the rate-limiting steps in the synthesis of melatonin at night (reviewed in ((Saha, Singh, & Gupta, 2018)), (Zhao et al., 2019)). As in vertebrates, melatonin is present in *Drosophila* (Finocchiaro, Callebert, Launay, & Jallon, 1988) with levels rising at night (Callebert, Jaunay, & Jallon, 1991; Hintermann et al., 1996), and *AANAT1* is thought to be the serotonin acetyltransferase used in its synthesis. Neither *AANAT1* mRNA or protein undergoes daily expression cycles in whole head extracts (Brodbeck et al., 1998), however melatonin synthesis might only occur in a small subset of cells. Interestingly, melatonin synthesis is lost in mutants of the circadian rhythm gene *period* (Callebert et al., 1991). In the Chinese Tasar Moth (*Antheraea pernyi*), the expression of an ortholog to *AANAT1* is under the control of clock genes in a subset of neurons, and RNAi against that *AANAT* lead to changes in photoperiodism and decreased melatonin (Mohamed et al., 2014). Similar to its role in dopamine metabolism, detailed analysis of *AANAT1* expression and its relation to serotonergic neurons may help identify locations of melatonin synthesis in the brain.

This project was the outcome of a research-based undergraduate course. Three students with no knowledge of *Drosophila* genetics or development mapped the *speck* mutation to the *AANAT1* gene within one semester. They acquired valuable skills in a multitude of areas including: science history, literature searches, genetics, molecular biology, photomicroscopy, writing, and scientific reasoning. While this project mapping historical mutants was successful, any number of similar projects could be designed where students work to solve problems and make discoveries. These research-based, problem solving courses are not just better for the students, but more enjoyable for the faculty (E. Spana, unpublished observation).

## Supporting information

Supplemental Movie 1: Oregon R

Supplemental Movie 2: speck-2 blistered-2

## ACKNOWLEDGEMENTS

Thanks to the Duke University Biology Department for course funding and Duke University Office of Undergraduate Research for research grants to ABA and KTE. Thanks to Jamie Roebuck of Model System Injections for the CRISPR injections and to Dylan Ryan for developing the method for the time lapse wing expansion videos. We would especially like to thank Daniel Chun, David Jung and Bo Sun for their work on mapping *curved*, and Zachary Powell and David Rothschild for their work on mapping *spread*. Thanks to Dan Kiehart (Duke University) and Teruyuki Niimi (Nagoya University) for fly stocks. We thank Don Fox (Duke University Medical Center), Danny Miller (University of Washington), Jonathan Massey (University of Michigan), Neal Silverman (University of Massachusetts Medical School) and Rebecca Spokony (Baruch College) for their helpful comments on this manuscript.

## MATERIALS AND METHODS

### *D. melanogaster* culture, handling, and crosses

All stocks were obtained from the Bloomington Drosophila Stock Center at the University of Indiana except for the UAS-Bm-aaNAT/TM3 (obtained from Teruyuki Niimi, Nagoya University) and the *y* w\** and *y* w*; bw^1^ speck^1^ zip^2^*/SM6a (obtained from Dan Kiehart, Duke University). Crosses were performed and maintained at 25°C on standard cornmeal/molasses/agar based fly media.

### Minos reversion crosses

Flies of *w^1118^; Mi*[*ET1}AANAT1^MB02738^* were crossed to *w^1118^; noc^Sco^*/SM6a, P{hsILMiT}2.4 and the resulting larvae were heat shocked daily for 1 hour in a 37° water bath over 3-4 days. Single males or two virgin females of *w^1118^; AANAT1^MB02738^*/ SM6a, P{hsILMiT}2.4 were crossed to *w^1118^*; *Df(2R)BSC356*/SM6a. Progeny males in the next generation that lacked the mini-*w^+^* eye color associated with P{hsILMiT} were scored for presence of GFP and any *speck* phenotype. Ten independent lines (6 *speck^+^*, 4 *speck^-^*) were balanced over CyO.

### Adult tissue mounts and photography

Anesthetized 2-5 or 7 day old male flies were photographed with a Nikon D300S camera mounted to a Leica MZ12 stereomicroscope using a ring light for illumination. For time-lapse movies, newly eclosed adults were affixed at the notum to a micro-injection needle which was mounted on an ∼1 cm cube of modeling clay. An image was taken every 5 seconds using Camera Control Pro 2 software (Nikon) for a total of 2500 images. Individual .jpg images were combined into an .mpg movie at 30 frames per second using Time Lapse Assembler software Version 1.5.3 (by Dan Bridges, 2012).

### Molecular Biology

Genomic DNA was purified from wild type and mutant flies using the Qiagen DNeasy Blood & Tissue Kit (#69504) and stored at −20°C after isolation. PCR amplification of long products was completed using Qiagen LongRange PCR Kit (#206401). Sequencing was performed by Eton Biosciences (Research Triangle Park, NC).

### Whole Genome Sequencing

Two libraries consisting of genomic DNA purified from males of the genotypes *IP3K2^wy-1^*; SM6a/+ and from *t^1^ v^1^ m^74f^ IP3K2^wy-74i^*; *speck^2^ bs^2^/+* were used as template for HiSeq 4000, 150bp, paired-end sequencing (BGI Tech). Over 36.6 million reads for each library produced ∼5.5 Gigabases of sequence for each. A 20 kb region spanning the *AANAT1* region was assembled and analyzed using CONSED, version 29.0 (Gordon & Green, 2013) and minimap2, version 2.17-r941 (Li, 2018) and visualized using Tablet, version 1.19 (Milne et al., 2012).

### CRISPR mutations

Guide RNAs corresponding to two regions of the *AANAT1* gene were designed according to the method described in (Gratz et al., 2013). Phosphorylated oligonucleotides were obtained (IDT DNA), annealed and ligated into a linearized, dephosphorylated pU6-BbsI-chiRNA plasmid. A mix of both plasmids corresponding to the left and right breakpoints were injected into M{vas-Cas9}ZH-2A, *w^1118^*/FM7c by Model System Injections (Durham, NC). Injected G0 adults were crossed singly to *w^1118^; Mi{ET1}AANAT1^MB02738^* and the resulting progeny were scored for a *speck* phenotype. Three adults showing a *speck* phenotype were obtained from 66 crosses and new *speck* alleles were balanced over CyO, homozygosed and molecularly characterized by PCR amplification and sequencing.

### List of Sequences to be deposited

*speck^G1^* breakpoint

*speck^T1^* Mi perfect revertant

*speck^31B^* breakpoint

*speck^35A^* breakpoint

*speck^47B^* breakpoint

SM6a genome

*speck^2^ bs^2^* genome

### Supplemental Movies

Supplemental Movie 1: Wing expansion and cuticle tanning in wild type Oregon R Supplemental Movie 2: Wing expansion and cuticle tanning in *speck^2^ bs^2^*

### Supplemental Table of Primer sequences

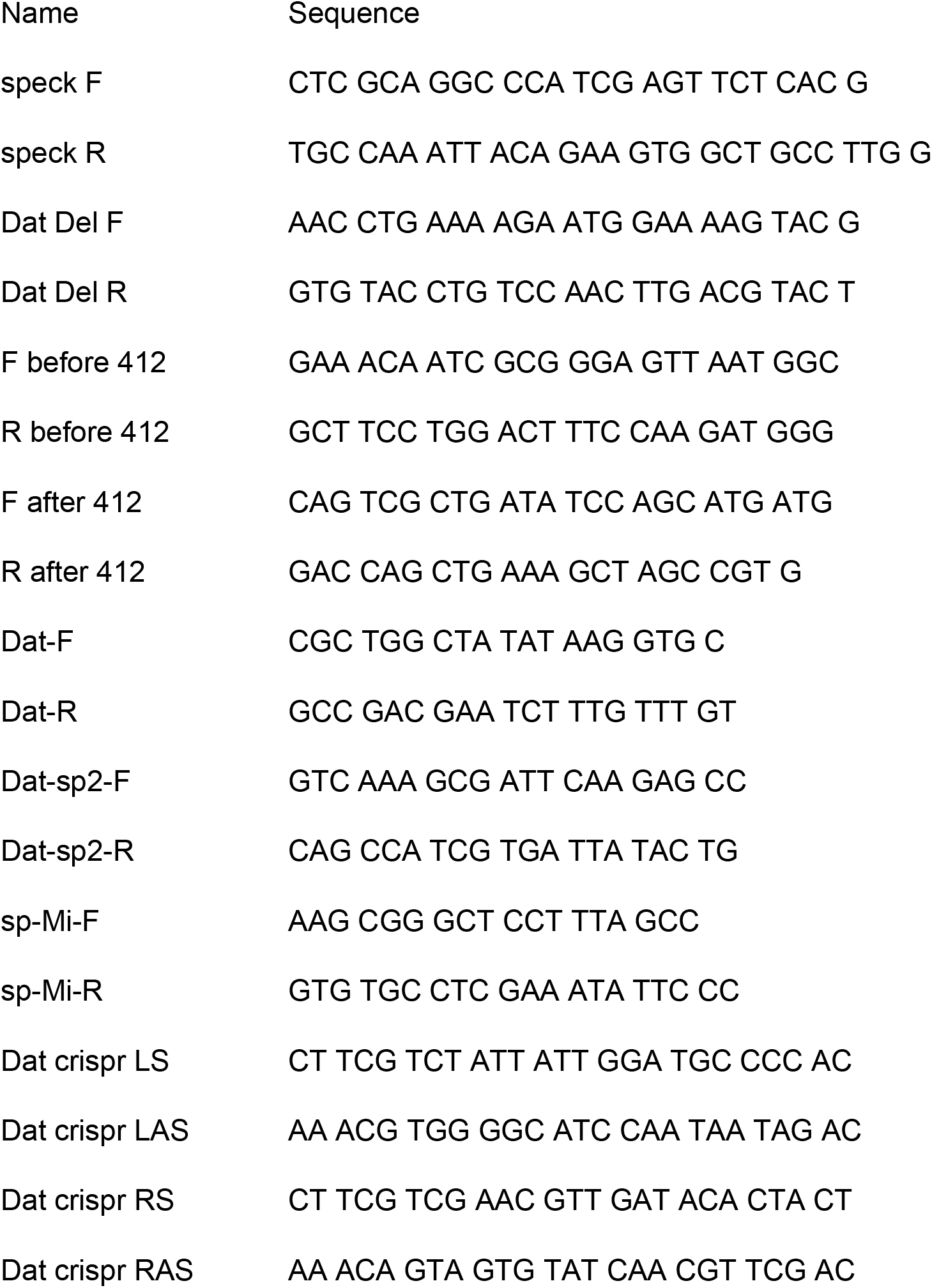

